# Genome-scale oscillations in DNA methylation during exit from pluripotency

**DOI:** 10.1101/338822

**Authors:** Steffen Rulands, Heather J Lee, Stephen J Clark, Christof Angermueller, Sébastien A Smallwood, Felix Krueger, Hisham Mohammed, Wendy Dean, Jennifer Nichols, Peter Rugg-Gunn, Gavin Kelsey, Oliver Stegle, Benjamin D Simons, Wolf Reik

## Abstract

Pluripotency is accompanied by the erasure of parental epigenetic memory with naïve pluripotent cells exhibiting global DNA hypomethylation both *in vitro* and *in vivo*. Exit from pluripotency and priming for differentiation into somatic lineages is associated with genome-wide *de novo* DNA methylation. We show that during this phase, coexpression of enzymes required for DNA methylation turnover, DNMT3s and TETs, promotes cell-to-cell variability in this epigenetic mark. Using a combination of single-cell sequencing and quantitative biophysical modelling, we show that this variability is associated with coherent, genome-scale, oscillations in DNA methylation with an amplitude dependent on CpG density. Analysis of parallel single-cell transcriptional and epigenetic profiling provides evidence for oscillatory dynamics both *in vitro* and *in vivo*. These observations provide fresh insights into the emergence of epigenetic heterogeneity during early embryo development, indicating that dynamic changes in DNA methylation might influence early cell fate decisions.

**Highlights:**

- Co-expression of DNMT3s and TETs drive genome-scale oscillations of DNA methylation
- Oscillation amplitude is greatest at a CpG density characteristic of enhancers
- Cell synchronisation reveals oscillation period and link with primary transcripts
- Multiomic single-cell profiling provides evidence for oscillatory dynamics *in vivo*

## Introduction

In mammalian embryonic development, the segregation of lineages giving rise to different somatic tissues is associated with large-scale changes in DNA methylation (5-methylcytosine). Following fertilisation, global loss of DNA methylation from both the maternal and paternal genomes is tightly linked with the acquisition of naïve pluripotency in the inner cell mass of the blastocyst (Lee et al., 2014). During the transition towards the primed pluripotent state of the epiblast, *de novo* methylation results in a global gain of this epigenetic mark (Auclair et al., 2014; Seisenberger et al., 2012; Smith et al., 2012; Wang et al., 2014). A similar event occurs *in vitro* during the transition from naïve to serum primed embryonic stem cells (ESCs) and then exit from pluripotency (Ficz et al., 2013; Habibi et al., 2013; Leitch et al., 2013; Takashima et al., 2014; von Meyenn et al., 2016). However, during priming in ESCs not only are the *de novo* methyltransferases DNMT3A and B dramatically upregulated but paradoxically the hydroxylases TET1 and 2 remain highly expressed too. This observation has given rise to the hypothesis that the system may be dynamic with turnover of DNA methylation to hydroxymethylation (5hmC), formylcytosine (5fC), carboxycytosine (5caC) through base excision repair or DNA replication back to unmethylated cytosine (Lee et al., 2014). This could potentially lead to heterogeneous epigenetic states between cells in a population, with functional consequences for gene expression and cell phenotype. DNA methylation and chromatin dynamics have been quantitatively modelled in various genomic contexts in bulk datasets and in exquisite detail at single loci of biological significance (Atlasi and Stunnenberg, 2017; Berry et al., 2017; Bintu et al., 2016; Haerter et al., 2014; Kyriakopoulos et al., 2017). However the availability recently of methylome information in single cells from single-cell whole genome bisulfite sequencing (scBS-seq, Farlik et al., 2015; Smallwood et al., 2014) provides an unprecedented opportunity for studying and modelling DNA methylation dynamics genome-wide in a population of cells undergoing a biological transition. Indeed scBS-seq has revealed profound methylation heterogeneity in ESCs particularly in enhancers (Farlik et al., 2015; Smallwood et al., 2014). Here we combine single-cell sequencing with biophysical modelling to study how DNA methylation heterogeneity and dynamics arise during the transition from naïve to primed pluripotency, and the exit from pluripotency *in vivo*. We find evidence for genome-scale oscillatory dynamics in these cells with a link to primary transcripts, suggesting that heterogeneity can be created by molecular processes on different scales.

## Results

### Heterogeneous methylation distributions in primed ESCs

To study DNA methylation heterogeneity during the phase of lineage priming we began by considering ESCs, which serve as a powerful *in vitro* model for cells transiting from naïve through primed pluripotency and into early cell fate decision-making (Kalkan et al., 2017). scBS-seq measurements have shown that methylation heterogeneity in ESCs is greatest at enhancer elements (e.g. H3K4me1 sites and low methylated regions (LMRs) (Stadler et al., 2011)) (Smallwood et al., 2014). By analysing scBS-seq data separately for naïve and serum conditions, we found that increased variance at H3K4me1 sites was specific to primed ESCs (Figures 1A and S1A), with cell averages varying between 17% and 86% (Figures 1B and 1C). (Note that, to avoid potential systematic variations in DNA methylation during the cell cycle, ESCs were isolated based on their DNA content and were predominantly positioned in the G0/G1 phase.) By contrast, naïve ESCs showed minimal cell-to-cell variability at H3K4me1 sites (Figures 1B, 1C and S1A). Although other genomic elements showed proportionately less variability, levels of DNA methylation at these sites were found to be tightly correlated with those at enhancer regions and highly coherent for CpG poor elements (Figures 1D and S1B). DNA methylation heterogeneity in enhancer regions therefore is a reflection of synchronized (coherent) changes that affect the DNA on a genome scale.

**Figure 1.**
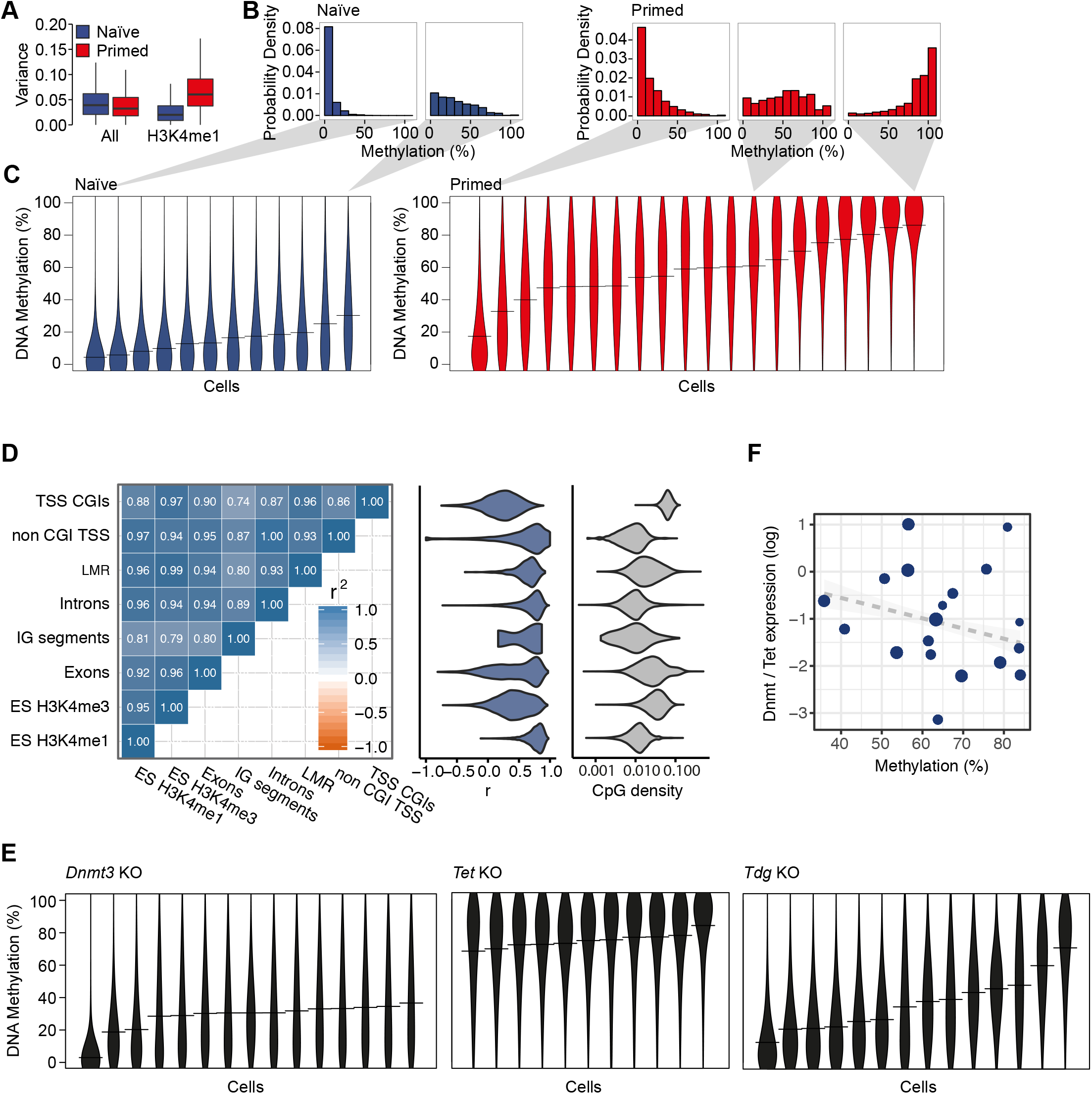
Correlated heterogeneity in DNA methylation. (A) DNA methylation variance in naïve and primed ESCs is compared for 3kb tiles over the whole genome (All), or for tiles overlapping H3K4me1 sites. (B) Histograms of DNA methylation at H3K4me1 sites for selected individual naïve and primed ESCs. (C) Violin plots of DNA methylation at H3K4me1 sites for individual cells from naïve and primed ESCs. (D) Left: Correlation of global average methylation across cells for different genomic features. Middle: Distribution of Pearson’s correlation coefficient between methylation levels at specific sites and global average H3K4me1 methylation. Right: Distributions of CpG densities as defined by the number of CpGs divided by the number of base pairs. (E) Violin plots of DNA methylation at H3K4me1 sites for individual Dnmt3, TET1-3 and TDG knock-out (KO) ESCs. (F) Scatter plot comparing H3K4me1 methylation and transcription of DNA methylation modification enzymes in scM&T-seq data from ‘More Pluripotent’ primed ESCs (see Figure S2C). The size of dots is proportional to global methylation coverage.

Methylation of cytosine residues is catalysed by the *de novo* DNA methyltransferase (DNMT3A/B) enzymes, while ten-eleven translocase (TET1/2/3) enzymes act in a multi-step process that can remove DNA methylation (Wu and Zhang, 2014). Importantly, primed ESCs express both *Dnmt3a/b* and *Tet1/2*, while naïve ESCs express *Dnmt3a/b* at much reduced levels (Figure S2A), raising the possibility that DNA methylation heterogeneity is dependent on this paradoxical co-expression (Lee et al., 2014). Consistently, we observed a loss of DNA methylation heterogeneity during differentiation to embryoid bodies, where *Tet* and *Dnmt3* are downregulated (Figures S2A and S2B). Analysis of parallel single-cell DNA methylome and transcriptome sequencing (scM&T-seq) data (Angermueller et al., 2016) showed that DNA methylation heterogeneity at H3K4me1 marks is largely confined to the most pluripotent sub-population, which express the highest levels of *Dnmt3a/b* and *Tet1/2* (Figure S2C). Furthermore, deletion of *Dnmt3a/b* resulted in homogeneously low DNA methylation levels, while loss of *Tet1-3* led to uniformly high DNA methylation (Figure 1E and S2D).

How does this strongly correlated DNA methylation heterogeneity arise during the transition from naïve to primed pluripotency? One possibility is that methylation differences between primed ESCs reflect slow dynamic changes in the expression of DNMT3A/B and TET1/2 arising, for example, through transcriptional state switching (Singer et al., 2014). However, although DNA methylation heterogeneity is dependent on the coexpression of genes that drive methylation and demethylation, analysis of scM&T-seq data (Angermueller et al., 2016) shows that global methylation levels (i.e. the genome-wide mean methylation rates) are largely independent of their (Figure 1F, R^2^=0.06). Moreover, DNA methylation is dynamic in primed conditions (Singer et al., 2014) and, as a system in steady-state, it follows that such dynamics must be recurrent. Such recurrent dynamics could be achieved by DNA methylation switching stochastically and reversibly between discrete levels or, alternatively, by continuously oscillating. Crucially, since we observe strong genome-wide coherence in DNA methylation levels (Fig. 1D), such recurrent changes in DNA methylation must be synchronized (i.e. coherent) across the genome.

### Modelling dynamics of DNA methylation

To assess whether DNA methylation turnover could give rise to oscillatory dynamics, we turned to a modelling approach. Importantly, our approach was constrained by the observed genome-wide coherence of DNA methylation levels, placing emphasis on finding a description based on collective degrees of freedom. We therefore set out to ask whether and how collective dynamics in DNA methylation can emerge despite the plethora of complex heterogeneities that can influence DNA methylation locally.

We began by considering the dynamics of a single CpG site, which can assume several different states including an unmodified cytosine (C), a methylated cytosine (5mC), a hydroxymethylated cytosine (5hmC), and other states. Notably, the biochemistry of DNA methylation turnover involves a cyclical process: The binding and action of DNMT3A/B drives conversion of C to 5mC (Baubec et al., 2015; Jia et al., 2007), while demethylation occurs through a long sequence of intermediary steps, each requiring the binding and release of enzymes, and ultimately the excision of intermediates and DNA repair or DNA replication (Figure 2A). Importantly, DNMT3A/B has been shown to bind cooperatively to the DNA (Emperle et al., 2014), implying that *de novo* methylation is autocatalytic. Meanwhile, the removal of DNA methylation effectively leads to a time delay, *Δt*, between the removal of the 5mC mark and the reestablishment of the unmodified cytosine. Given the known coupling between histone modifications, chromatin remodelling and DNA methylation (Du et al., 2015; Iurlaro et al., 2017), it is likely that these different levels of regulation contribute to the nonlinear feedback of DNA methylation on itself. Mathematically, we reasoned that the time evolution of C and 5mC concentrations, *c*(*t*) and *m*(*t*), averaged across the genome, can therefore be captured by the minimal coupled set of rate equations (Supplementary Theory),

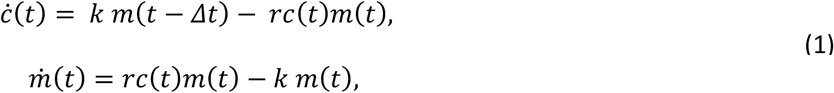

with *k* and *r* defining effective chemical conversion rates from C and 5mC, respectively. If the time delay *Δt* is sufficiently long, *de novo* methylation of initially hypomethylated genomic regions will result in rapid depletion of the pool of unmodified cytosines, which is then filled again due to the delayed conversion of 5mC to C. This can then lead to sustained oscillations in the levels of C and 5mC through a mechanism termed a Hopf bifurcation (Figure 2B and 2C). Importantly, although we do not know the effective conversion rates *k* and *r* in living cells, the fact that the model predicts coherent oscillations for low values of the dimensionless parameter *kΔt* suggests that coherent oscillations can occur under biologically relevant conditions. Indeed, distributions of methylation rates obtained from stochastic simulation of the dynamics resemble closely the experimental distributions obtained by scBS-seq (Figure 2D, Supplementary Movie 1 and Supplementary Note).

**Figure 2.**
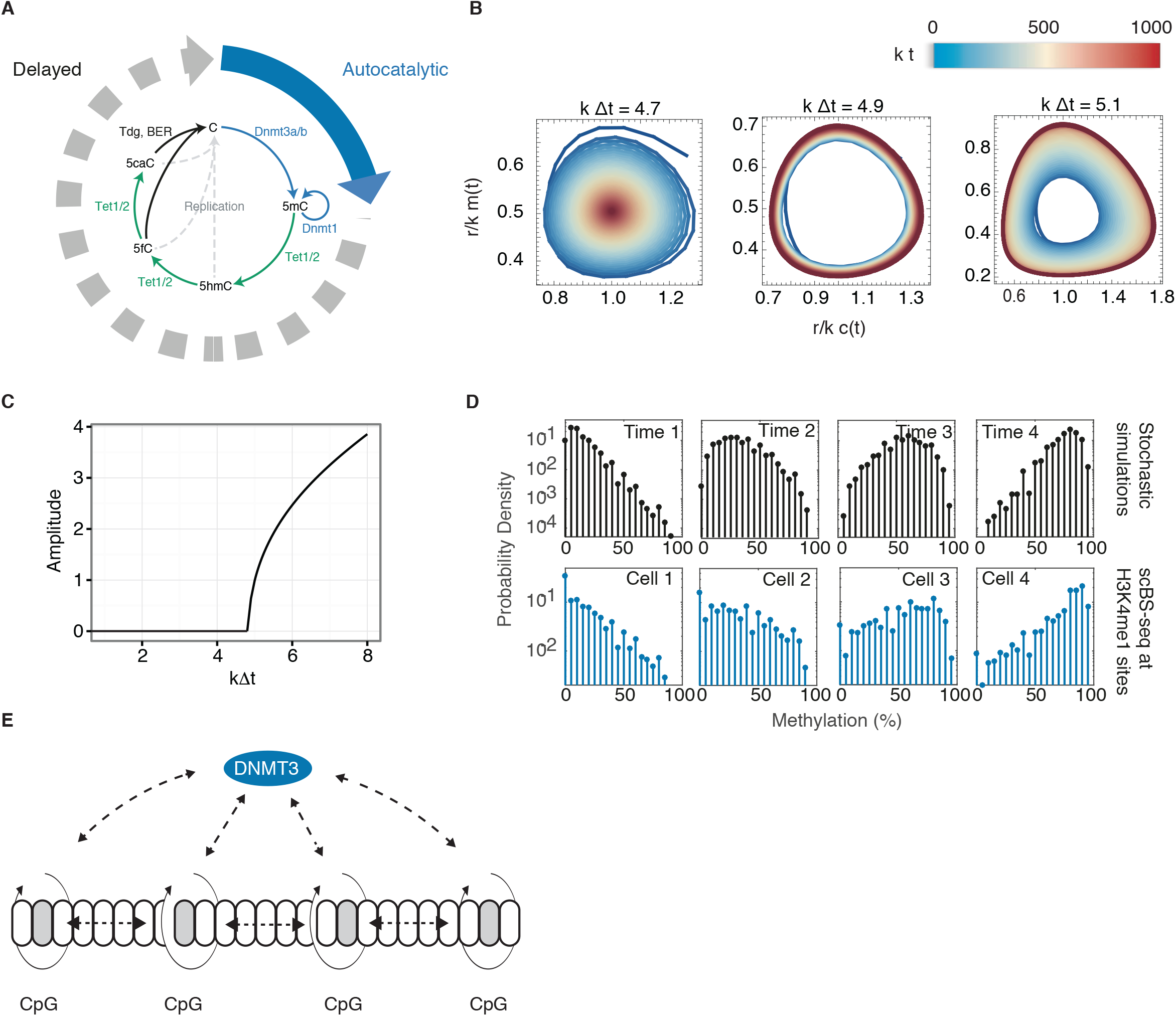
Biophysical modelling of DNA methylation turnover predicts global oscillations in DNA methylation. (A) Schematic summarising the biochemical processes involved in the turnover of cytosine modifications and a simple biophysical model comprising autocatalytic *de novo* methylation and time-delayed de-methylation. (B) Numerical solution of Equation (1) for dimensionless concentrations of (un-) methylatied CpGs for various values of the dimensionless delay time *kΔt*. Color denotes time, such that early times are blue, intermediary time yellow and late times are red. (C) Amplitude of oscillations as a function of the dimensionless time delay. (D) Top: Distributions of methylation rates at H3K4me1 sites as obtained from stochastic simulations. Panels show different time points of the simulations. Bottom: Exemplary distributions of DNA methylation rates at H3K4me1 sites in different cells obtained from scBS-seq experiments. (E) Schematic summarising global and local modes of coupling of CpGs via DNMT3a/b binding.

Although this simple model captures the essence of how global oscillations may emerge from the biochemistry of DNA methylation turnover, its validity relies implicitly on some mechanism by which information on methylation levels is transported throughout the genome. How can such collective behaviour, as indicated by the experimental data, arise given the known heterogeneity of local factors influencing DNA methylation? To answer this question, we turned to a more *ab initio* modelling approach, considering the stochastic dynamics of individual CpG sites which, according to the biochemistry of DNA methylation turnover, cycle through multiple chemical states stochastically with a locus-specific rate. We hypothesised that coherent collective dynamics can emerge as a result of the autocatalytic binding of DNMT3A/B enzymes. These enzymes can methylate multiple neighbouring CpGs at the same time, leading to their effective short-range coupling (Haerter et al., 2014). At the same time, DNMT3A/B enzymes preferentially bind to 5mC, which represses active degradation of these enzymes and thereby leads to global positive feedback on DNA methylation (Figure 2E). We took both local and global feedback to be locally heterogeneous mirroring local variations in enzyme binding affinity (conferred e.g. by different chromatin contexts).

To investigate whether locally heterogeneous interactions can give rise to global oscillations of DNA methylation, we successively coarse grained the stochastic dynamics starting from CpG dense regions and progressing to CpG poor regions (using an approach known as strong coupling renormalisation). For further details of the method and its current application, see Supplementary Theory. During this process, neighbouring blocks of CpGs become increasingly uncoupled, such that the coarse grained local phase dynamics is effectively described by a heterogeneous Kuramoto model,

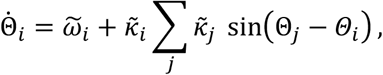

with continuous phases, *Θ_i_*, and effective intrinsic frequencies 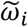 and couplings 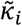. The dynamics is therefore described effectively by a set of globally coupled driven oscillators. Just as our phenomenological model, the heterogeneous Kuramoto model exhibits collective oscillations through a Hopf bifurcation if the average coupling through DNMT3A/B binding is sufficiently strong,

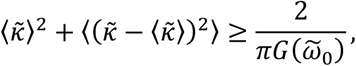

where 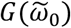 of the probability of the most abundant oscillator frequency.

This result suggests that, first, coherent oscillations can occur due to local and global feedback by DNMT3A/B enzymes, and that this effect is enhanced by heterogeneity in DNMT3A/B binding affinities. Second, if DNMT3A/B mediate global DNA methylation oscillations we expect the oscillation amplitude in a given genomic region to be proportional to the rate of DNMT3A/B binding in this region. Third, due to the cyclic nature of DNA methylation turnover, the local frequency should, at least transiently, be inversely proportional to the rate of DNMT3A/B binding. Importantly, it is these predictions that we use to challenge the basis of the model below using the experimental findings.

### Evidence for rapid DNA methylation oscillations upon serum priming

To obtain more direct evidence for genome-scale DNA methylation oscillations, and to test these model predictions, we next considered an *in vitro* “2i release” model in which cells were transferred from naïve “2i” to primed “serum” culture conditions and bulk cell samples were collected for BS-seq over a subsequent time-course (Figure 3A). As naïve ESCs show homogeneously low DNA methylation levels (Figure 1C), we reasoned that transfer from naïve to primed conditions might synchronise their entry into an oscillatory phase, allowing direct evidence for oscillations to be acquired from population-based measurements. Strikingly, we detected evidence for rapid oscillations in the mean methylation rate over H3K4me1 marks, with a period of approximately 2-3h (Figures 3A and S3A-C). Oscillations in global methylation were also observed in other genomic contexts, such as CpG-poor promoters and exons (Figures 3B, 3C and S3C). Spectral analysis confirmed enriched oscillations in H3K4me1 (p=0.05) and H3K27ac elements (p=0.007), as well as exons (p=2e-4), introns (p=8e-7), promoters (p=0.01) and genome-wide (p=1e-51) (Figures 3E and S3C). In agreement with the model, from the spectral analysis, we found that the period of oscillations during the release from 2i differed between genomic elements (Figure 3E and S3C), being longer at specific enhancer regions known to repress DNMT3 binding than elsewhere. Indeed, this initial variability in periodicity indicates that oscillations in DNA methylation are not driven extrinsically by a global (genetic) oscillator, such as Hes1 (Kobayashi and Kageyama, 2011), which would lead to the same single harmonic at all regions of the genome.

**Figure 3.**
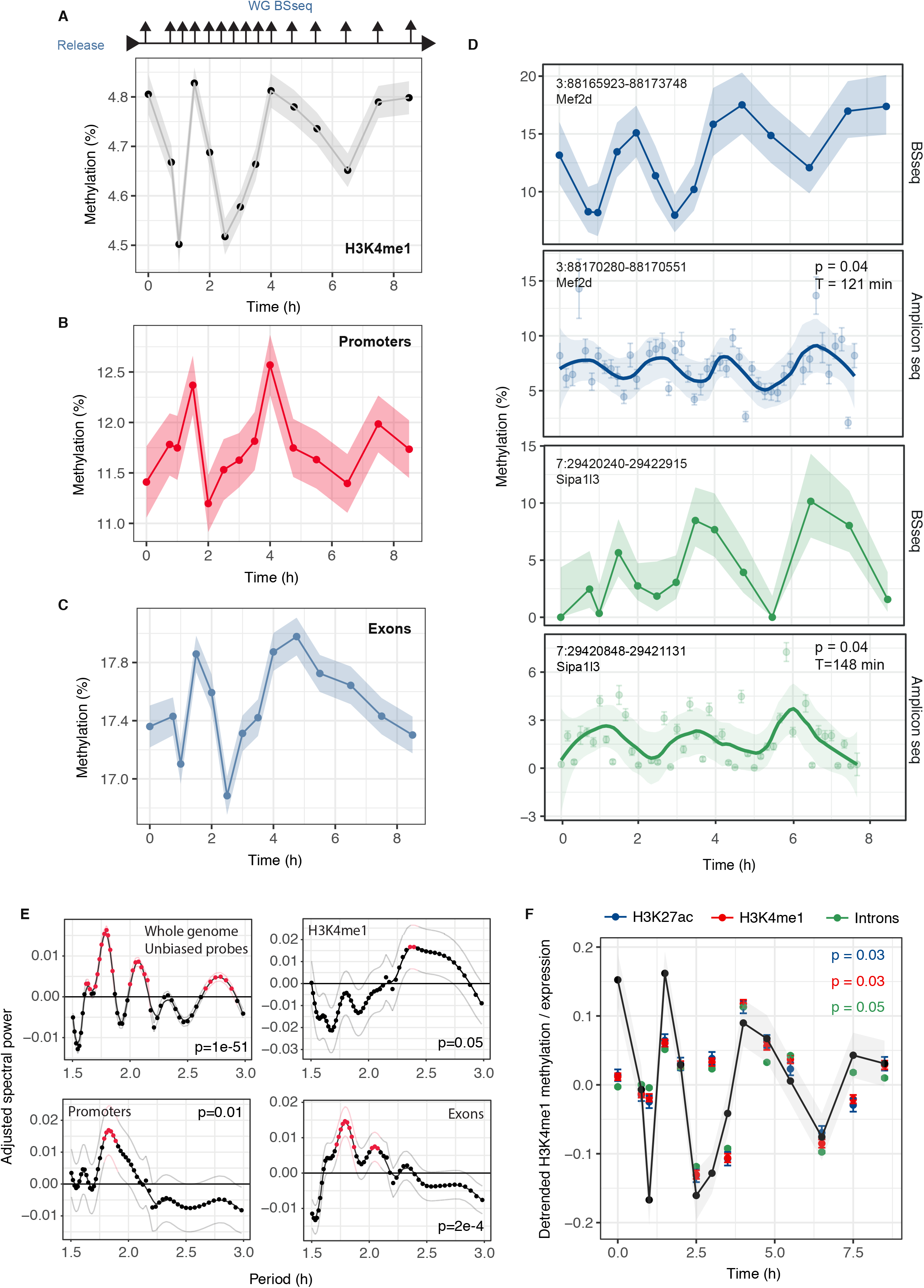
Oscillatory dynamics of DNA methylation during transition from naïve to primed pluripotency *in vitro*. (A) Average DNA methylation at H3K4me1 sites over the time course. For the average, we took into account 50% of enhancers with the highest coverage depth over the time course. (B) Average methylation at promoter regions and (C) exons. (D) Methylation levels at exemplary enhancer elements as measured by BS-seq (top) and AmpBS-seq (bottom). Lomb-Scargle spectral analysis was performed on the AmpBS-seq time course. (E) Average spectral densities for different genomic features (see also Figure S3C). Red dots denote significant enrichment of a given period (p<0.05). Thin lines denote standard error. (F) Comparison between the DNA methylation time course in H3K4me1 regions (see also Figure 3A) and average log-expression in different genomic contexts after removal of slow trends (see Materials and Methods). Shaded regions and error bars in (A-C),(D), (F) represent standard error.

The amplitude of genome-wide oscillations was seemingly modest; however, global averages in bulk measurements represent only the residual signal after averaging over many noisy elements, and may be confounded by cell-to-cell variability in the timing of DNMT3A/B up-regulation upon priming such that these measurements can only provide a lower bound for the amplitude. Indeed, oscillations with substantially greater amplitude were found upon inspection of specific H3K4me1 sites (Figures 3E and S3D). Oscillations were more subtle in a repeat experiment and could not be rigorously resolved with the coverage depth available in whole-genome bisulfite sequencing. However, using amplicon bisulfite sequencing (AmpBS-seq) to target 14 loci (Table S2) at H3K4me1 sites that showed evidence of oscillatory dynamics in the initial 2i release experiment, we confirmed oscillations at 4 of these 14 loci upon 2i release using spectral analysis (Figure 3D and S3D), while no oscillations were observed in cells that remained in 2i media. Furthermore, when considering a larger set of 35 loci by AmpBS-seq, spectral analysis revealed significantly enriched oscillations compared to the control experiment (p=0.05, Fisher’s test).

To explore the potential functional impact of DNA methylation oscillations on transcription, we performed an RNA sequencing time course of the same samples after release from 2i conditions. While results for exonic reads were ambiguous, we found significant correlations in enhancer and intronic reads with global DNA methylation levels (Figure 3F). This difference in the strength of correlation in exonic and non-exonic reads could be a reflection of the shorter half-lives of these primary transcripts compared to mature mRNA.

### Oscillations are CpG density-dependent

To further probe the mechanistic basis of DNA methylation oscillations and challenge the model predictions, we returned to the initial 2i release experiment to investigate whether oscillations were equally prevalent across the genome or preferentially enhanced in specific genomic elements. The locally averaged distance between neighbouring CpGs, or its inverse, the CpG density, defines a natural scale in the context of DNA methylation (Lövkvist et al., 2016). We therefore tiled the genome into windows of variable length, but constant local sequencing coverage (50 informative CpGs), to account for varying CpG coverage (see Figure 4A for details of the approach). For each window, we then determined the CpG density and the amplitude of oscillation upon 2i release. We found that the amplitude diverged at a characteristic value of the CpG density of around 2.5%, while oscillations were largely suppressed at CpG-rich regions (Figure 4B).

**Figure 4.**
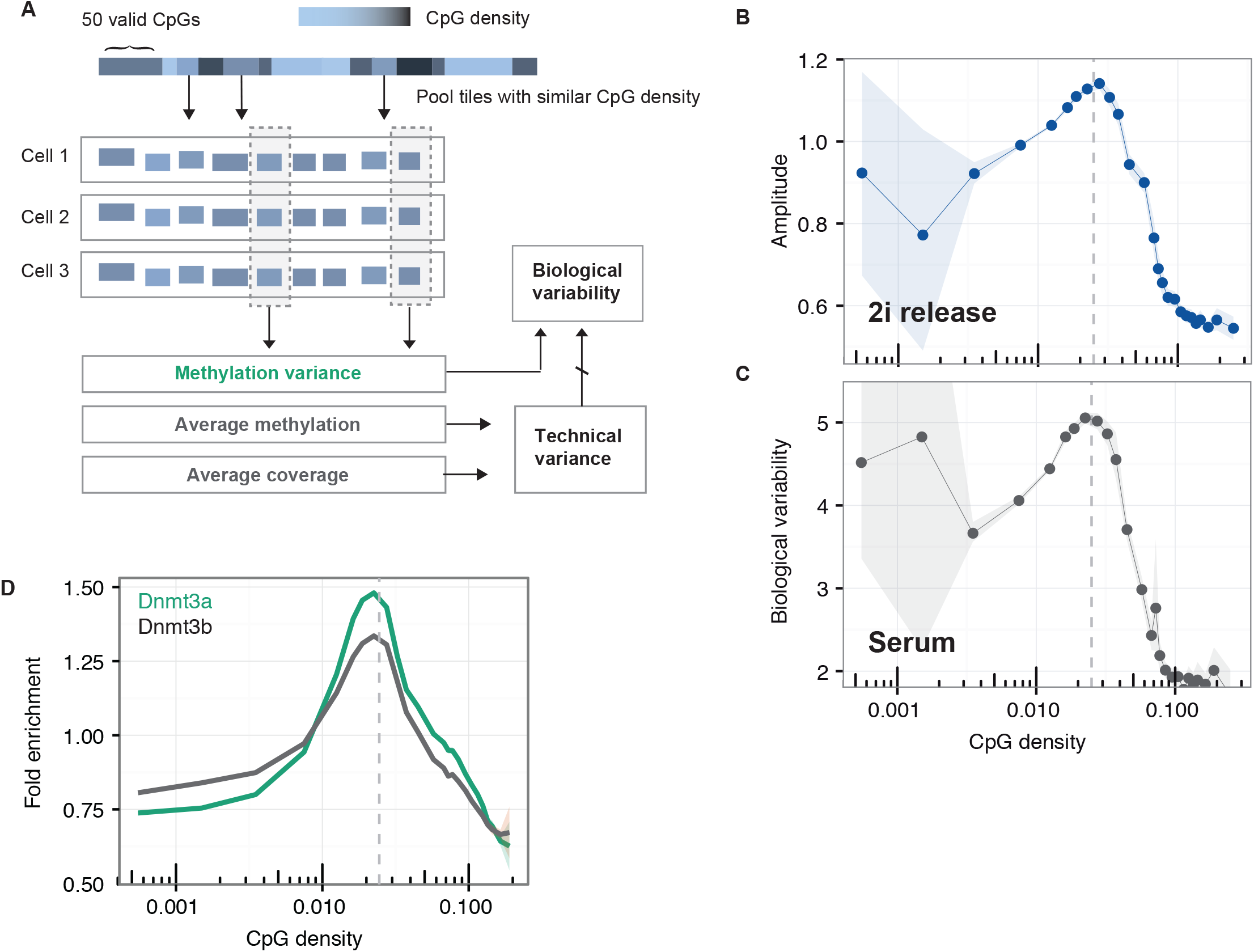
CpG Density is a key parameter defining oscillatory dynamics. (A) Illustration of the analysis of CpG-density dependent methylation: We segmented the genome into tiles of 50 consecutive informative CpGs (unbiased probes). We then grouped regions with similar CpG density and calculated the average methylation level for a given CpG density, the standard deviation between cells, and the average coverage in given region. Biological variability was then calculated as the biological variability over technical variance as an indicator of the amplitude of oscillation (B) Amplitude of oscillations of DNA methylation following transfer from naïve (2i) to primed conditions, as a function of CpG density. (C) An analogous analysis reveals biological variability as a function of CpG density in a long-term culture of primed ESCs. (D) Fold enrichment over input of Dnmt3a/b binding as a function of CpG density. Chip-seq data were analysed similarly to (Baubec et al., 2015). We tiled the genome into 1kbp tiles with an overlap of 500bp and added 8 pseudo counts per element.

Based on this observation, we returned to the scBS-seq data for primed ESCs in steady state and calculated how much cell-to-cell variability in DNA methylation exceeds that expected from technical noise. To estimate biological variability for a given locus, and to take into account confounding factors to methylation variance, we followed previous work and considered the ratio of methylation variance across cells and the technical variance expected for a given combination of mean methylation and coverage (see Methods). Notably, we found the same CpG density-dependent divergence as for the amplitude of oscillations after 2i release (Figure 4C), consistent with our hypothesis that methylation heterogeneity in primed ESCs derives from oscillatory dynamics. Moreover, the divergence in the strength of oscillations coincides with the measured CpG density-dependence of DNMT3A/B binding affinity (Figure 4D) suggesting that, in agreement with the model, coherence is mediated through DNMT3A/B binding.

### Evidence for coherent oscillations of DNA methylation in vivo

Noting that the transcriptional and epigenetic changes that occur in ESCs following their transfer from 2i to serum conditions resemble those seen *in vivo* during the exit from pluripotency (Kalkan et al., 2017), we then questioned whether oscillatory DNA methylation dynamics can be observed in the embryo. Indeed, during this transition (E4.5 to E5.5 epiblast), there is a substantial increase of *Dnmt3a* and *b* transcript levels while *Tet1* remains highly expressed (Boroviak et al., 2015; Mohammed et al., 2017), suggesting that co-expression of these enzymes could drive oscillations *in vivo*. We therefore analysed parallel scM&T sequencing of epiblast cells at E4.5, E5.5 and E6.5 (Argelaguet et al., unpublished). Once again, we observed cell-to-cell variability in the levels of DNA methylation at primed ESC enhancer sites (Figures 5A and S4A). At E4.5 and E6.5, global DNA methylation correlates with transcriptional changes associated respectively with early and late lineage priming (Argelaguet et al., unpublished). However, at E5.5, as in primed ESCs, global methylation levels at enhancers were largely independent of *Dnmt3* and *Tet* expression levels in the same cell (Figure 5B, *R*^2^ = 0.12) and the transcriptome did not show any early signs of lineage priming (Mohammed et al., 2017; Peng et al., 2016). Moreover, at this time point, DNA methylation heterogeneity was also independent of any genes that vary spatially across the embryo at E6.5 (Supplementary Theory) (Scialdone et al., 2016).

**Figure 5.**
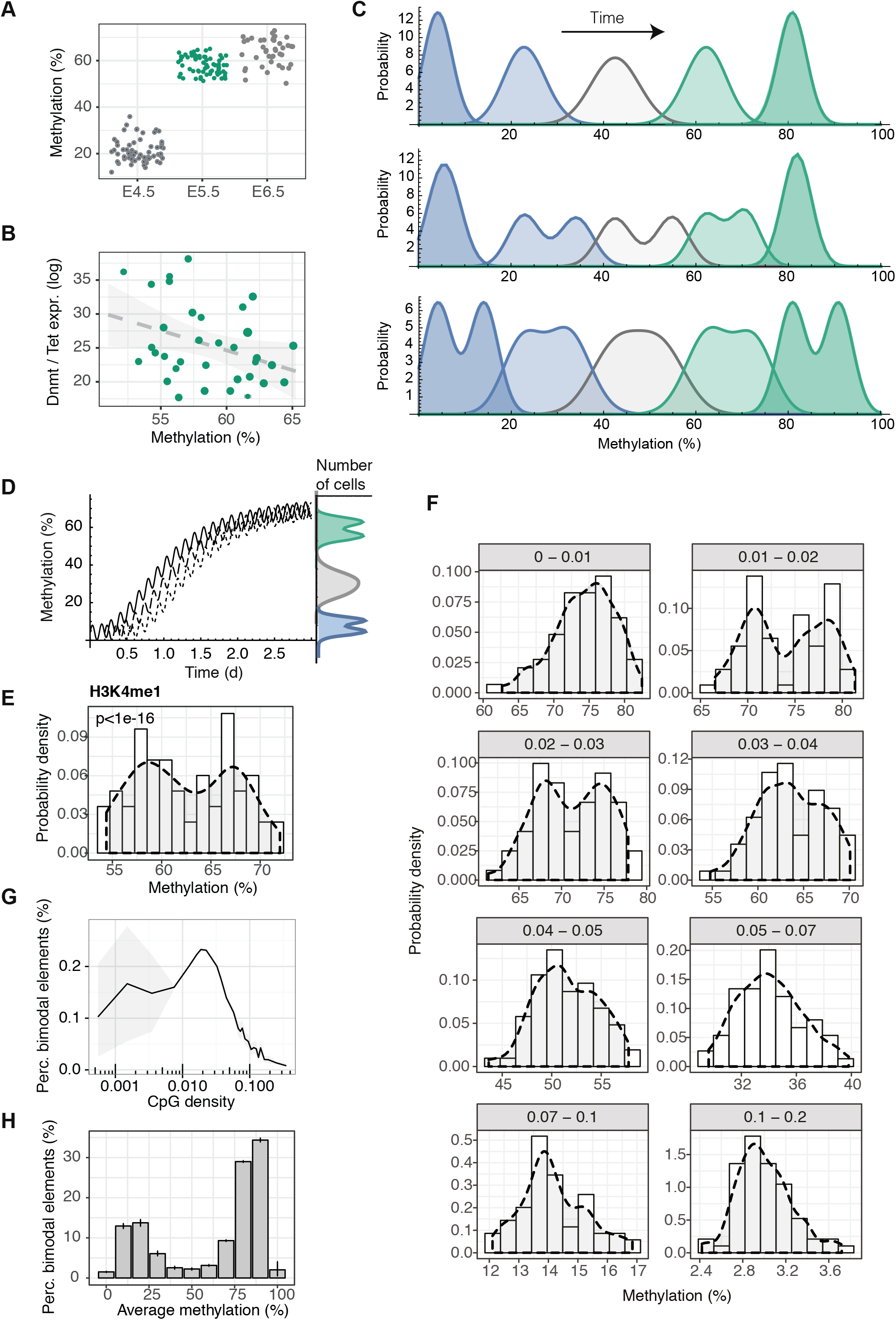
scMɖT-seq reveals evidence for oscillatory DNA methylation *in vivo*. (A) Global DNA methylation levels around H3K4me1 sites of individual cells at three stages during early mouse embryo development. (B) Global DNA methylation levels around H3K4me1 sites versus expression levels of genes that positively influence methylation over expression levels of genes that drive demethylation. (C) Predictions of the distribution of global DNA methylation levels during the process of global *de novo* methylation for various time points in different scenarios (for details, see Supplementary Theory). Top: If oscillations are absent and the *de novo* methylation is initiated at time points following a unimodal distribution the distribution of global DNA methylation levels remains unimodal at all times (average methylation levels). Middle: If *de novo* methylation is initiated at early time points following a bimodal distribution (early lineage segregation) the distribution of global DNA methylation is bimodal at intermediary times (average methylation levels). Bottom: If global *de novo* methylation is superimposed with oscillatory dynamics we expect bimodality at early and late times (low and high average global methylation levels), but not at intermediary times (intermediary average global methylation levels). (D) Schematic illustrating the specific patterns *in vivo* for oscillating global DNA methylation. In contrast to alternative scenarios, biophysical modelling predicts a bimodal distribution of average methylation levels at early and late stages of global *de novo* methylation if methylation dynamics has an oscillatory component (for details, see Supplementary Theory). (E) Probability density of global DNA methylation levels around H3K4me1 sites reveals evidence for bimodality. (F) Probability distributions (bars: from histograms, shaded areas: density estimation) of DNA methylation levels taking into account regions with different ranges of CpG densities. For this analysis the genome was tiled into windows of 50 consecutive informative CpGs. (G) Fraction of unbiased probes (100 valid CpGs length) that show statistically significant patterns of bimodality (dip-test, p<0.05) as a function of CpG density. (H) Fraction of unbiased probes that show statistically significant bimodality as a function of their average methylation level across cells.

Based on these observations, we hypothesised that the heterogeneity of DNA methylation at E5.5 is a consequence of *de novo* methylation or oscillatory turnover. But how can oscillatory dynamics be identified from a purely static measure, such as that provided by single-cell sequencing? To address this question, we first sought to identify statistical patterns in DNA methylation that are specific for oscillatory DNA methylation. We reasoned that static measurements of a population of cells exhibiting oscillations around the same centre point would, with higher probability, reflect cells at the extrema of the oscillation than at intermediary values. Therefore, if the progressive increase in *de novo* methylation is superimposed with oscillatory dynamics, the distribution of the average levels of DNA methylation would become bimodal both at the onset of this transition and when DNA methylation has reached saturation levels (Figure 5C,D and Supplementary Theory). By contrast, during the transient phase of increasing global DNA methylation levels, cell-to-cell variability in this process would overshadow this bimodal signature resulting in a unimodal distribution. Alternative hypotheses, such as variability in the timing of entry into the primed phase, would ultimately lead to unimodal distributions of global DNA methylation levels during the transition period with the peak tracking the increase in the average level of DNA methylation (Supplemental Theory).

At E5.5, when global DNA methylation levels were near maximal level, we found that the distribution of methylation was indeed bimodal in enhancer regions (p<1e-16) and other genomic contexts (Figures 5E,F and S4B), consistent with oscillatory DNA methylation. To further challenge the association of bimodality with oscillatory dynamics, we tiled the genome into coverage-based windows and used a statistical (dip) test to assess whether DNA methylation in a given window is bimodally distributed between cells. In excellent agreement with the divergence of the oscillation amplitude after 2i release and in serum conditions, the bimodal signature was strongest for elements with approximately 2.5% CpG density (Figures 5F,G and S4B). Further, independent of CpG density, we compared genomic regions at different stages of the *de novo* methylation process. In agreement with oscillatory dynamics (Figure 5D), and independent of CpG density, bimodality was only pronounced at hypomethylated or hypermethylated regions, but not at regions with intermediary methylation level (Figure 5H). Notably, although our analysis does not rule out early lineage commitment through DNA methylation heterogeneity, such a scenario cannot explain the observed CpG density-dependence of bimodality or the depletion of the bimodal pattern at regions with intermediary DNA methylation levels.

Finally, to obtain more direct evidence for DNA methylation oscillations *in vivo*, we sought to resolve oscillations by ordering cells according to an estimate of their “developmental age”, i.e. the time since the initial upregulation of the *Dnmt3* genes. To this end, we noted that CpG-rich regions do not show pronounced oscillations *in vitro*, and DNA methylation levels in these regions rise monotonically between E4.5 and E6.5 (Figure 6A). We therefore used global methylation levels in regions with a CpG density of between 10% and 15% to define a methylation “pseudotime” for individual cells (Supplementary Theory). Then, charting the average DNA methylation levels of genomic elements with CpG densities for which oscillations are expected to be most pronounced (i.e. between 2 and 3%) against pseudo-time, we found evidence of coherent oscillatory patterns, which were then confirmed using spectral analysis (p=6e-4, Figures 6B and 6C). In common with the findings of the 2i release experiment, this oscillatory pattern was strongest at approximately 2.5% CpG density (Figure S4B).

**Figure 6.**
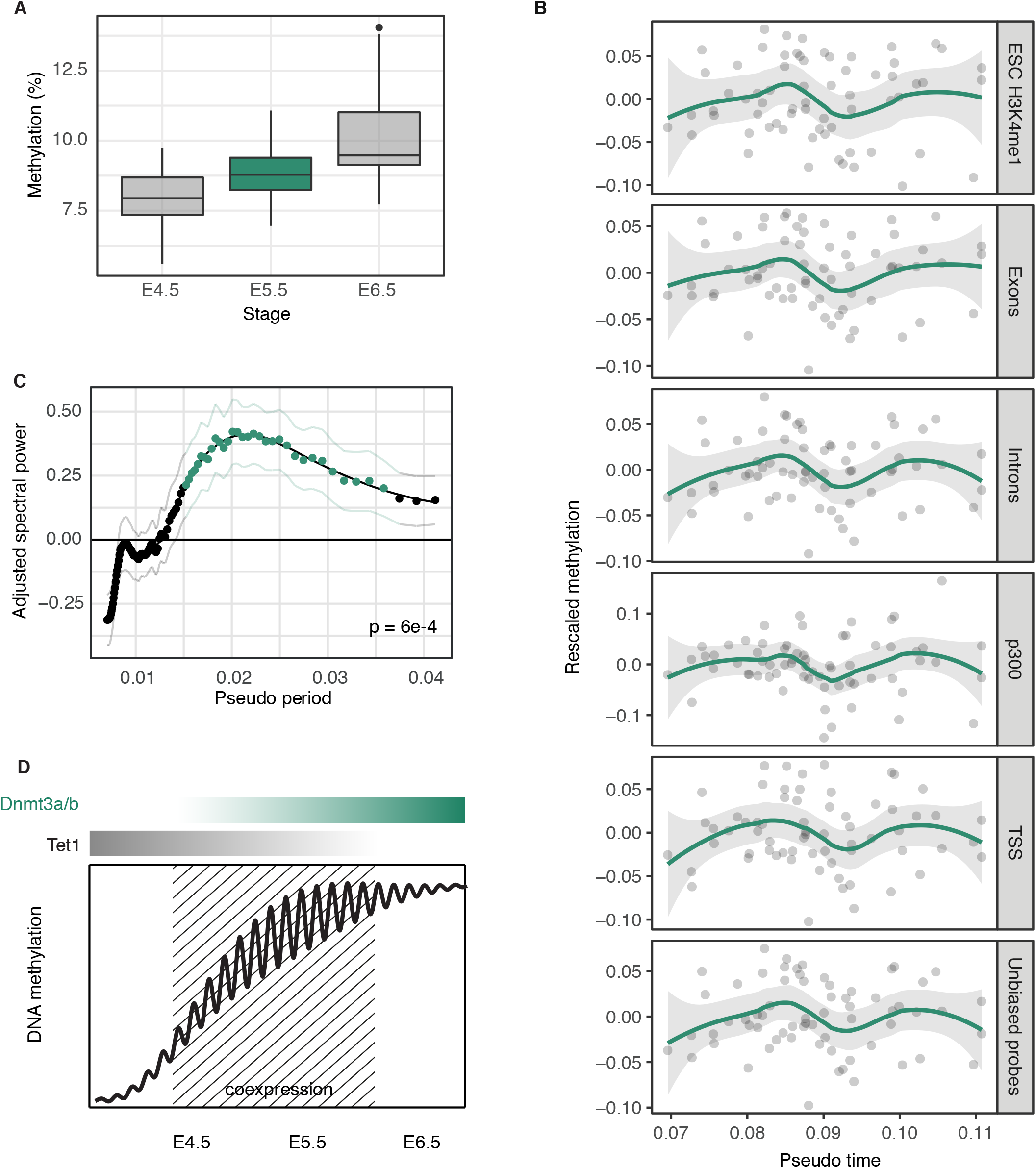
Pseudo-time analysis provides independent evidence for *in vivo* oscillations of DNA methylation. (A) Box plots of average DNA methylation levels of individual cells at three stages during early mouse embryo development acquired from genomic regions with CpG densities between 10 and 15%. (B) Using the average methylation levels from (A) as a measure of the “developmental time” of a given cell, DNA methylation levels in different contexts show parallel non-monotonic dynamics. (C) Average spectral densities for the whole genome. Red dots denote significant enrichment of a given period (p<0.05). Thin lines denote standard error. (D) Summary schematic depicting the trend for DNA methylation levels at sites of intermediate CpG density during the exit from pluripotency. As levels of DNA methylation rise during this phase, co-expression of Dnmt3s and Tets promote intermittent genome scale oscillations of DNA methylation.

## Discussion

Transcriptional and epigenetic heterogeneity between cells is thought to be important for cell fate decision making during development (Torres-Padilla and Chambers, 2014), but the underlying molecular mechanisms are largely unknown. Using single-cell sequencing technologies, we have systematically investigated the dynamics of DNA methylation heterogeneity during the exit from pluripotency and priming for differentiation. By combining biophysical modelling and single-cell sequencing, we have revealed evidence for genome-scale oscillations in DNA methylation during the exit from pluripotency (Figure 6D). Mechanistically these oscillations appear to be driven by cooperative binding properties of the DNMT3 enzymes, which makes low CpG density sequences, including enhancers, a particular target. Signatures of oscillations are also present in embryos exiting pluripotency *in vivo* suggesting the possibility that they may contribute to cell fate decisions in embryogenesis.

Based on the DNA modification cycle, mathematical modelling predicts the emergence of genome-scale oscillations in DNA methylation in cells where both DNMT3 and TET enzymes are expressed. These conditions arise naturally during priming of ESCs and in epiblast cells *in vivo*, when DNMT3A/B levels increase strongly in cells already expressing TET1/2, before overt differentiation leads to the down-regulation of both DNMT3 and TET enzymes. The oscillation state is hence an intermediate between the naïve hypomethylated state and the differentiated state in which the majority of the enhancer sites in the genome are methylated. By synchronising cells in the naïve state and then measuring DNA methylation genome-wide at closely spaced time points upon serum priming, we were able to record robust oscillations in DNA methylation, which occurred with a period of approximately 2-3 hours. Given the multi-step cycle of cytosine modification turnover, these oscillations are remarkably fast. However, yet more rapid oscillations in DNA methylation (with a period of approximately 1.7 hours) have been observed in breast cancer cells at the *pS2* promoter upon transcriptional activation (Kangaspeska et al., 2008; Metivier et al., 2008).

DNA methylation oscillations in primed ESCs are more rapid than, and therefore must be autonomous of, the cell cycle and the rate of switching between transcriptional states. Of course, longer period transcriptional switching of the *Tet* genes may influence the oscillation dynamics at the population level (see below). However, it is notable that our model can yield oscillations at constant levels of DNMT3a/b and TET1/2 if one considers nonlinearities, such as autocatalytic and processive *de novo* methylation, where established 5mC marks catalyse further *de novo* methylation (Baubec et al., 2015). In accordance with the model, we found that oscillations stalled at a low point of DNA methylation upon removal of *Dnmt3a/b* and at a high point upon *Tet1/2* depletion.

Our initial focus was on enhancer methylation, the sites of greatest heterogeneity in primed ESCs. This is consistent with LMRs and H3K4me1 sites being targeted by hydroxymethylation in ESCs and being the most methylation-variable sequences between tissues upon differentiation *in vivo* (Booth et al., 2012; Feldmann et al., 2013; Hon et al., 2013; Hon et al., 2014; Huang et al., 2014; Lu et al., 2014; Stadler et al., 2011; Ziller et al., 2013). At first sight, the amplitude and the genome-scale synchronisation of oscillations might seem inconsistent with limited accessibility of DNA in condensed chromatin. However, transcription factor binding to enhancers and promoters disrupts the local nucleosome structure rendering chromatin more accessible, as reflected in DNAseI hypersensitivity and ATAC-seq assays. We found that many regions of the genome participate in oscillatory methylation in a manner that is dependent on CpG density.

With a period of 2-3 hours, DNA methylation is surprisingly dynamic, changing at a rate much faster than cell division or transcriptional state switching. Parallel BS-seq and RNA-seq sequencing during 2i release suggests that oscillations in DNA methylation are correlated with changes in primary transcripts, pointing to a potential functional role. Intriguingly, in parallel with the current study, genome-scale oscillations with approximately the same period of 2-3 hours have been observed through studies of nascent transcription at intronic sites in mESCs in serum conditions (L. Cai et al, manuscript in revision at *Cell*). Through alterations in DNA binding affinities for the transcriptional machinery mediated by changes in DNA methylation, these findings point at periodic changes in “Waddington’s epigenetic landscape” that occur on similar or faster time-scales than those of cell lineage decisions. Future developments in single-cell multi-omics and the manipulation of epigenetic states *in vivo* will determine whether and how oscillations in DNA methylation play an instructive role in promoting transcriptional heterogeneity with consequences for symmetry breaking and lineage priming.

## Acknowledgements

*Dnmt3a/b* knock-out ESCs were the generous gift of Dr Tatyana Nesterova; *Tet1-3* knock-out ESCs were the generous gift of Dr Guoliang Xu; *Tdg* knock-out ESCs were the generous gift of Dr Primo Schär. We thank the Welcome Trust Sanger Institute sequencing pipeline team for assistance with Illumina sequencing. We thank Dr John Marioni for critical reading and discussion. W.R. is supported by BBSRC, Wellcome Trust, and EU. B.D.S. acknowledges the support of the Wellcome Trust (grant number 098357/Z/12/Z). H.J.L. is supported by EU NoE EpiGeneSys. G.K. is supported by BBSRC, MRC, and EU. O.S. is supported by EMBL, Wellcome Trust, EU.

## Author Contributions

H.J.L., S.R., B.D.S. and W.R. conceived the project and acquired funding. H.J.L., S.J.C. and S.A.S. performed experiments. S.R. developed theory and performed modeling. S.R. and C.A. performed statistical analysis. F.K. processed and managed sequencing data. G.K. and O.S. supervised technical aspects of the project. H.J.L., S.R., S.J.C., B.D.S and W.R. interpreted results and drafted the manuscript. All authors edited and approved the final manuscript.

## Data Deposition Statement

The accession number for sequencing datasets reported in this paper will be available on request. Published data sets used in analysis are listed in Table S4.

